# Large-Scale Quantitative Cross-Linking and Mass Spectrometry Provides New Insight on Protein Conformational Plasticity within Organelles, Cells, and Tissues

**DOI:** 10.1101/2024.11.14.623288

**Authors:** Andrew Keller, Anna Bakhtina, James E. Bruce

## Abstract

Many proteins can exist in multiple conformational states *in vivo* to achieve distinct functional roles. These states include alternative conformations, variable PTMs, and association with interacting protein, nucleotide, and ligand partners. Quantitative chemical cross-linking of live cells, organelles, or tissues together with mass spectrometry provides the relative abundance of cross-link levels formed in two or more compared samples, which depends both on the relative levels of existent protein conformational states in the compared samples as well as the relative likelihood of the cross-link originating from each. Because cross-link conformational state preferences can vary widely, one expects intra-protein cross-link levels from proteins with high conformational plasticity to display divergent quantitation among samples with differing conformational ensembles. Here we use the large volume of quantitative cross-linking data available on the public XLinkDB database to cluster intra-protein cross-links according to their quantitation in many diverse compared samples to provide the first widescale glimpse of cross-links grouped according to the protein conformational state(s) from which they predominantly originate. We further demonstrate how cluster cross-links can be aligned with any protein structure to assess the likelihood that they were derived from it.

## Introduction

Within living systems, molecular function required to support life is achieved through a complex and dynamic set of intra- and inter-protein interactions defined by all conformations and protein-protein interactions existent within the cellular interactome. Functional and interactome regulation is mediated by modulation of proteome levels but is also achieved through changes in conformations and interactions that occur independent of protein abundance level changes. While quantitation of proteome levels with mass spectrometry has become a routine and indispensable tool in efforts to map cellular function, visualization of conformational and interaction changes inside cells to better understand how these mediate functional adaptation has remained challenging.

Chemical cross-linking and mass spectrometry constitute an emerging approach, yielding both qualitative and quantitative information on the cellular interactome that can improve understanding of biological function^1–10^. Many proteins can exist in multiple conformational states with alternative conformations^11,12^, variable modifications^13,14^, and a diversity of associated interacting protein^15^, nucleotide^16^, and ligand^17^ partners, all of which serve to shape protein conformational plasticity. The probability of forming and observing a given intra-protein cross-linked product is affected by proximity of its reactive amino acid sites due to the finite cross-linker physical length, as well as the accessibility and reactivity of those sites and the occurrence of PTMs on all residues of the cross-linked peptides^18^. Individual cross-links may preferentially originate from an alternative protein conformation^19^, from the presence or absence of a PTM^20^, or from the presence or absence of an interacting partner^21^. Here the term ‘cross-link’ is used to refer to cross-linked peptide pairs. The relative intra-protein cross-link abundance levels in phenotypic^19,22^, pharmacological^23,24^ or other cellular comparisons are thus reflective of all protein conformational states from which those cross links can originate and provide insight into many facets of protein conformational plasticity.

For the hypothetical comparative case of a protein with a single conformation that also has no changes in modification or interacting protein levels, one would expect all of its intra-protein cross-links to have levels that reflect the abundance of the sole protein conformational state, and thus to have similar quantitation in all compared samples equal to the relative change in protein levels. In the case of a protein with two alternative conformations, A and B, the relative abundance of any intra-protein cross-link that can originate with equal likelihood from either conformation would reflect the relative protein levels, as above. In contrast, the levels of intra-protein cross-links that are exclusive to or more probable to form in conformation A or B, such as those with reactive sites less than the maximum cross-linker length only in A or B, would reflect the relative abundance of those individual conformations, respectively. As a result, these three groups of cross-links should each display differing quantitation across compared samples that have varying relative levels of the two conformations (Figure 1).

**Figure 1.**
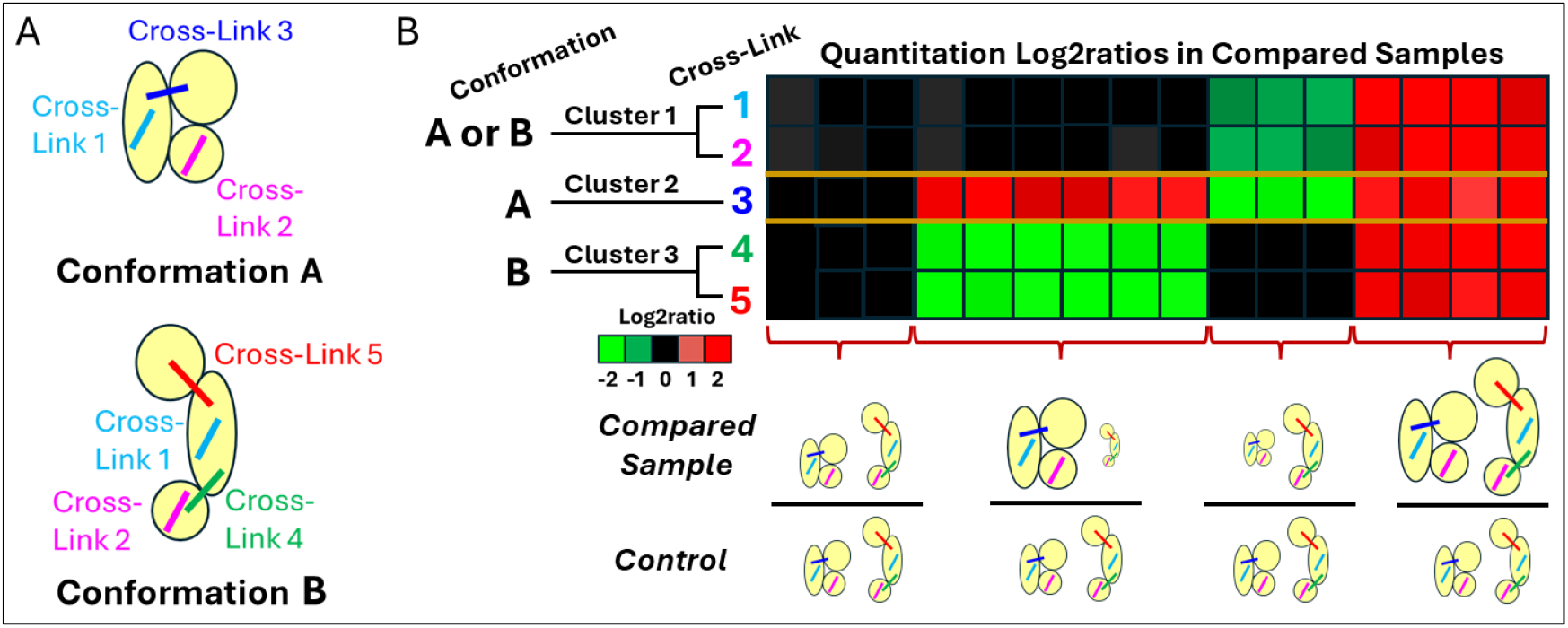
Intra-protein cross-link quantitation of protein with two conformations, A and B. A. Intra-protein cross-links #1 and #2 are obtained with equal likelihood from both protein conformations, #3 only from conformation A, and #4 and #5 only from conformation B. B. Heatmap of quantitation of cross-links in compared samples reveals 3 clusters of cross-links with abundances reflecting levels of total protein, conformation A, and conformation B, respectively. Below the heatmap are shown abundances of the two conformations in compared samples with respect to control, indicated by cartoon size, giving rise to observed quantitation patterns. The first has samples with no difference in levels of either conformation, the second has a sample with increased conformation A, decreased conformation B, and no overall protein level difference, the third has reduced levels of only conformation A, and the fourth, increased levels of both conformations.

In the case of a protein often modified with a PTM at a particular residue, the abundance of cross-links containing that residue with or without the PTM would reflect only the levels of the modified or unmodified protein, respectively, and display quantitation different from those cross-linked products that are insensitive to the presence of the PTM across compared samples with varying PTM levels. Similarly, in the case of a protein with an interacting partner, the abundance of intra-protein cross-links predominantly obtained from the dissociated proteins, such as those that can form at the inter-protein interface, would reflect the levels only of the dissociated protein and display quantitation different from intra-links that are insensitive to the protein association across compared samples with varying degrees of interaction with the partner protein. In the case of a multimeric association, a presumed intra-protein cross-link may also be derived from two protein molecules.

Because quantitation of intra-protein cross-link levels in compared samples can reflect changes in abundance of particular protein conformational states, the comparison and clustering of intra-protein quantitative cross-link levels across many and diverse sample comparisons can uniquely improve understanding of conformational plasticity that exists within cells. Of interest is differentiation of proteins that exhibit just a single cluster of intra-protein cross-links with similar quantitation, as would be expected if changes are due to abundance level changes of a lone conformational state, from those that exhibit multiple clusters indicative of adaptation involving conformational plasticity, where cross-links in each cluster have distinct quantitation patterns reflecting differing likelihoods of originating from multiple protein conformational states such as alternative conformations, PTMs, and interacting partners. Because of these multifaceted influences on cross-link levels, interpreting quantitative cross-link data is often challenging. Nevertheless, quantitation clustering gives a direct indication of the variety of encountered protein conformational states.

The growing number of quantitative XL-MS (qXL-MS) datasets on the public cross-link database XLinkDB^25,26^ with diversity of compared samples enables investigation of qXL-MS data and its potential to reveal protein conformational plasticity in living systems. Here we describe the analysis steps of intra-protein cross-link quantitation clustering and discuss its implications. We show how the clustering can help characterize the conformational state(s) likely giving rise to intra-protein cross-links in various clusters. Importantly, these efforts highlight unique value that quantitative *in vivo* cross-linking data hold for improved structural analysis of proteins in living systems. These efforts also underscore the potential that further increased accumulation of such data across the growing qXL-MS community could have for systems-level quantitative structural biology. The clustering results are maintained as a community resource on XLinkDB.

## Methods

All intra-protein cross-links on XLinkDB originating from a *H. sapiens* or *M. musculus* protein that was quantified with iqPIR^27^ with respect to 10 or more sample comparisons were included in the clustering analysis. Each pair of cross-links originating from a particular protein was assessed for having similar quantitation according to the vector distance of their quantitation log2ratios in all sample comparisons (10 or more) in which they were both quantified with high confidence, having a 95% confidence ≤ 0.5 and 6 or more contributing quantified ions. The statistical significance of the distance was determined by generating null distributions of 1000 vector distances between quantitation of both compared cross-links and those of randomly selected log2ratio values from each included compared sample. A z-score of the cross-link pair quantitation distance with respect to each null distribution was calculated, and from each, a corresponding p-value. The maximum value of the two calculated p-values was used to ensure rigor. A low p-value indicates similar quantitation by virtue of a small vector distance of quantitation between the two cross-links that is unlikely due to chance.

Hierarchical clustering was performed by iteratively calculating distances between clusters as the average vector distance p-value of all pairs of cross-links in the two clusters, where the average was calculated as the exponentiated average of log p-values. The pair of clusters with the smallest average p-value were merged together provided that p-value was not greater than 0.01, or in a final step to merge remaining singleton clusters, 0.1. This lenient maximum p-value is aimed at clustering together cross-links with similar but not necessarily identical preferences for protein conformational states. Furthermore, we expect many cross-links with similar overall quantitation to differ in a small number of sample ratios that contain differing levels of less commonly encountered conformational states.

Clustering results of these proteins are available to the research community at the Protein Plasticity site of XLinkDB (https://xlinkdb.gs.washington.edu/xlinkdb/index.php) with columns indicating the number of clusters, number of clusters with two or more cross-links, number of cross-links, and an indicator of higher quality clustering whereby all cluster members have very similar quantitation patterns yet different from those of other clusters (see below). One can click on any protein to view its clusters in HTML showing cross-links of clusters along with heatmaps of their quantitation log2ratios in compared samples. The similarity of quantitation among cross-links within a cluster and between cross-links of different clusters is calculated as the average pairwise cross-link -10*log vector distance p-value. Higher confidence is given to clusters that each have a high calculated similarity of quantitation among members yet have low calculated similarity with quantitation of members of other clusters. This measure is indicated in the clustering results HTML page for each analyzed protein. Higher quality clustering is assessed as within cluster calculated similarity of quantitation being greater than or equal to 85 and between cluster calculated similarity of quantitation, less than 35 or less than 0.65 times the maximum calculated similarity of quantitation within the clusters.

## Results and Conclusions

XLinkDB now features quantitation clustering that groups intra-protein cross-links according to the protein conformational state(s) from which they most likely originate. Of 23,337 *H. sapiens* and *M. musculus* non-redundant intra-protein cross-links confidently quantified with iqPIR in one or more of 1,152 compared samples, 9,375 were quantified in 10 or more compared samples and subjected to clustering on the basis of their log2ratios. This resulted in derived quantitation clusters for 167 *H. sapiens* and 104 *M. musculus* proteins, exhibited online at https://xlinkdb.gs.washington.edu/xlinkdb/intraproteinQuantClusters.php (see a tutorial on viewing clustering results in **Supporting Information Video S1**). One can view heatmaps of the quantitation of cross-links in each cluster among the compared samples and view those cross-links in the context of any specified structure to help assess the likelihood that they originated from it. Cross-links with a Jwalk^28^ solvent accessible surface distance (SASD) within the 51 Å empirical maximum span of the iqPIR cross-linker^29^, and dead-end (DE) peptides with an attached lysine DSSP accessibility^30,31^ of at least 50 (Figure S1), are inferred to be consistent with the structure, and thus possibly derived from it. These data were acquired from a wide variety of quantified compared samples including those subjected to heat shock, heart failure, and aging. The specific details of the samples are not important, but the greater the variety of perturbations explored, the greater possibility there is to detect clustered cross-links originating from different protein conformational states.

The numbers of intra-protein cross-link quantitation clusters obtained for the analyzed proteins are shown in Figure 2A. The great majority of proteins, 80%, have more than a single cluster indicating multiple conformational states present in the compared samples. It is likely that more clusters will be observed with the addition of new quantified cross-links and/or compared sample conditions. If one approximates that cross-links are binary, either originating or not from any conformational state, then the number of possible quantitation clusters grows exponentially with the number of states, equal to one less than 2 raised to the power of the number of conformational states since each cross-link must originate from at least one state. Because it is unlikely to quantify cross-links with all possible conformational state preferences, and because states with low relative abundance in samples may not detectably affect cross-link quantitation, the number of observed clusters will likely be smaller. At most one cluster of any protein contains cross-links equally likely to originate from all its conformational states, and hence with levels reflecting total protein abundance in all compared samples, including those with differing relative levels of the conformational states.

**Figure 2.**
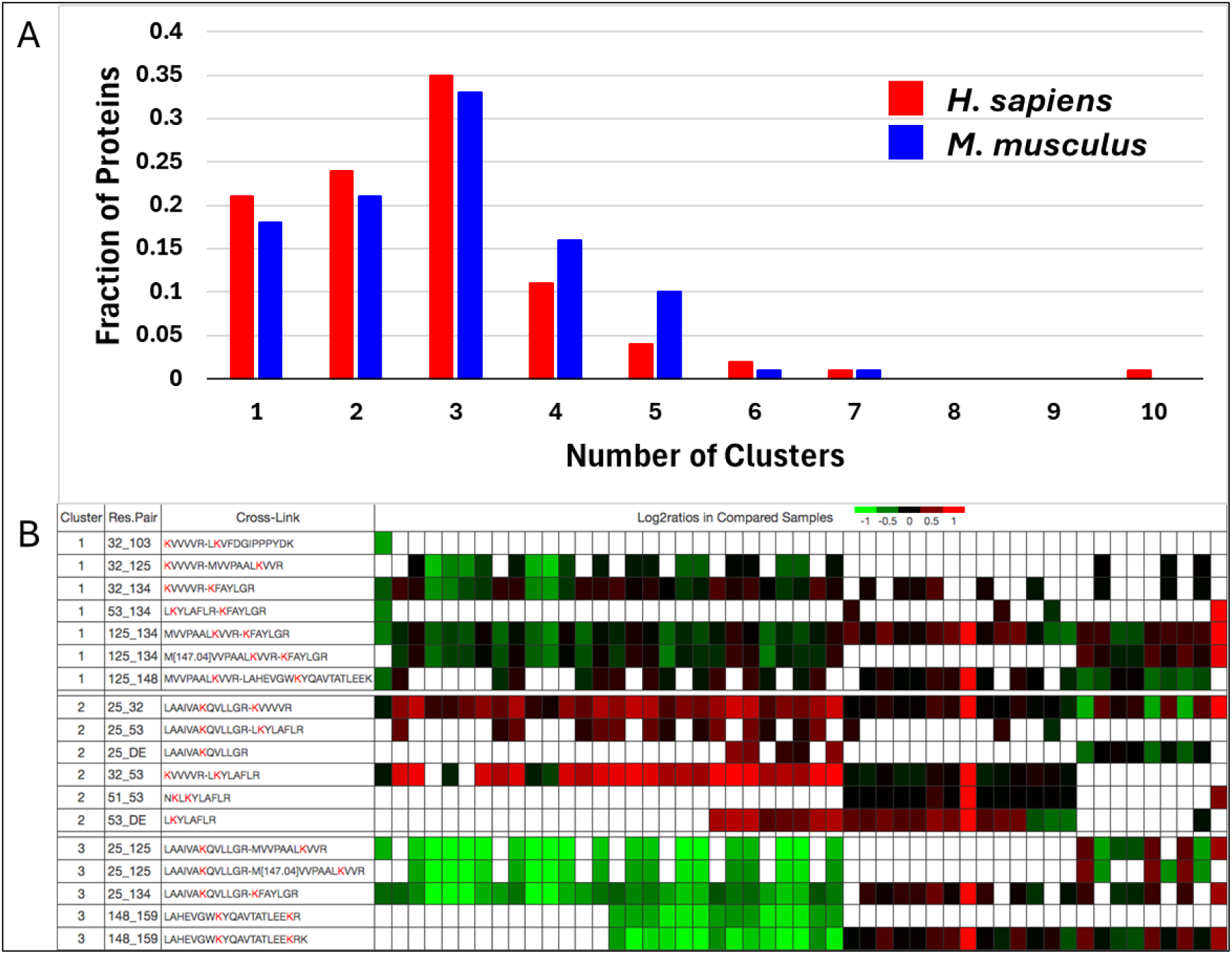
Intra-protein cross-link quantitation clusters. A. Distribution of the number of clusters obtained for *H. sapiens* and *M. musculus* proteins based on the quantitation of corresponding intra-protein cross-links. B. Heatmap showing distinct patterns in three clusters of the relative abundance of *H. sapiens* 60S ribosomal protein L13a intra-protein cross-links in compared samples. Log2ratios are indicated by color ranging from bright green (−1) to black (0) to bright red (+1) with missing values in white. A residue pair ending in _DE indicates a DE peptide.

Figure 2B shows distinct patterns of quantitation among compared samples displayed by intra-protein cross-links of RL13A_HUMAN, 60S ribosomal protein L13a, in its three major derived clusters. The quantitation patterns among cross-links within a cluster are similar to one another yet different from those of cross-links of other clusters, reflected by average calculated similarity of quantitation within and between clusters of 91 and 16, respectively (see Methods). Each cluster quantitation pattern reflects differing sensitivity of cross-link levels to two or more protein conformational states with altered relative abundance levels in some compared samples; no pattern can simply reflect levels of a lone conformational state. A component of the ribosome, this protein is expected to be present in multiple conformations and associated with multiple interacting partners.

Protein ADT2_HUMAN, ADP/ATP translocase 2, is a transporter known to have at least two distinct conformations, the m-state conformation open to the mitochondrial matrix and the c-state conformation open to the intermembrane space^32^. A third open channel conformation has also been proposed for stressed mitochondria^33^ and qXL-MS data was used with Alphafold2^34^ to produce an open model consistent with the proposed open state^19^. This protein has three quantitation clusters of cross-links. The first and second include cross-links found to be compatible with the m- and c-state conformations, respectively, as discussed below (Figure 3A).

**Figure 3.**
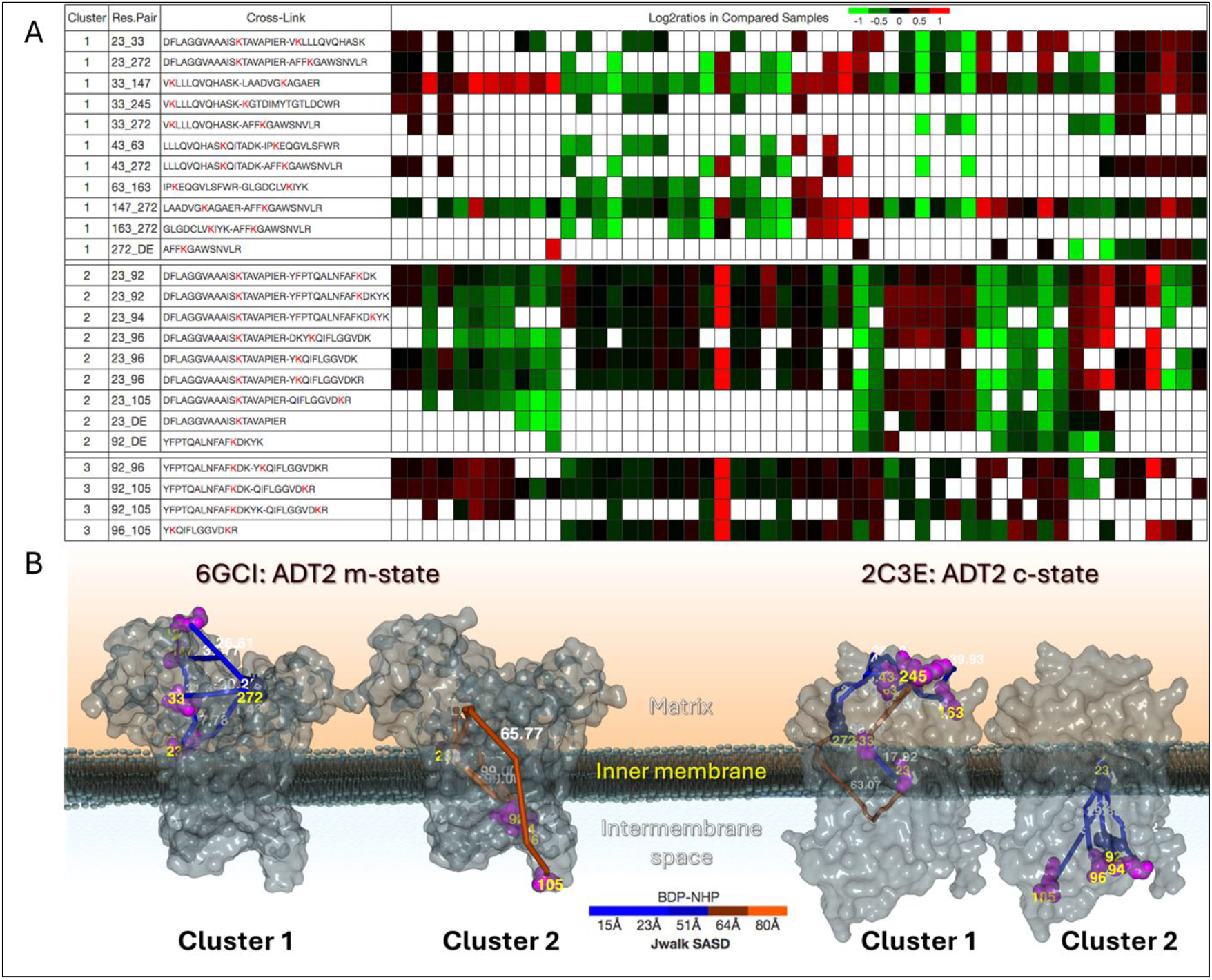
Three intra-protein cross-link quantitation clusters of ADT2_HUMAN. A. Quantitation of intra-protein cross-links of clusters in compared samples. Log2ratios are indicated by color ranging from bright green (−1) to black (0) to bright red (+1) with missing values in white. A residue pair ending in _DE indicates a DE peptide. B. Consistency of cross-links in clusters 1 and 2 with m-state structure 6GCI (channel open to the matrix) and c-state structure 2C3E (channel open to the intermembrane space). Cross-linked lysine residues are colored purple and their spans colored by Jwalk SASD consistency ranging from blue (consistent with cross-linker length) to black (borderline) to orange (in excess of cross-linker length). Cross-links in the first cluster are presumed to preferentially originate from the m-state, and those of the second cluster, from the c-state, conformations of ADT2_HUMAN.

According to SASDs of the cross-links and accessibilities of the DE peptide attached lysine residues in m-state structure 6GCI and c-state structure 2C3E, four of the 11 cross-links (including one DE peptide) in the first cluster, spanning residues 23_272, 33_147, 33_245, and 33_272, are inconsistent with 2C3E but consistent with 6GCI and are thus m-state specific (Figure 3B). Cross-links spanning residue pairs 147_272 and 23_33 have been reported to originate primarily from the m-state despite having Jwalk SASDs consistent with both state conformations^18,35^. Since a third open channel conformation has been proposed, some quantitation differences among the 11 cross-links may be due to differential likelihood of originating from that third conformation. Nevertheless, the clustering of quantitation data supports the notion that all cross-links in the cluster are indeed primarily m-state specific, including five not previously characterized as such, spanning residues 43_63, 43_272, 63_163, 162_272 and the DE peptide at residue 272. In contrast to the first cluster, all 9 of the cross-links (including two DE peptides) in the second cluster are inconsistent with structure 6GCI but consistent with 2C3E and are thus c-state specific. The four cross-links in a third cluster spanning residues 92_96, 92_105, and 96_105, are consistent with both structures and may be equally likely to originate from any conformation. Thus, the abundance of cross-links in the first, second, and third clusters are inferred to reflect levels of the m-state, c-state, and both state conformations, respectively.

In human cells, ADT exists in multiple isoforms, including ADT3_HUMAN, that have similar biological functions. ADT2_HUMAN and ADT3_HUMAN have cross-links spanning many of the same residue pairs, yet which are distinguishable due to sequence differences elsewhere in the cross-linked peptides, the only exceptions being a cross-link spanning residue 33_272 and a DE peptide at residue 92. Interestingly, the intra-protein cross-link quantitation clusters of ADT3_HUMAN match well with those of ADT2_HUMAN (Figure S2), demonstrating the ability of clustering to identify preferential protein conformational states giving rise to cross-links across related proteins. Of the cross-links observed in both isoforms spanning common residue pairs, the only clustering differences are two DE peptides that are placed with another DE peptide into a separate cluster of ADT3_HUMAN, and a lone cross-link spanning residues 92_105 that was included in the first cluster of ADT3_HUMAN alongside m-state specific cross-links. It is possible that with additional compared samples and quantified cross-links, that cross-link will be clustered separately as in the case of ADT2_HUMAN. Importantly, the clustering of ADT2_HUMAN intra-protein cross-link quantitation was also found to be similar to that of its *M. musculus* homologue of 99% sequence identity, ADT2_MOUSE, with 13 of their 15 common cross-linked residue pairs identically grouped (Figure S2). Only the cross-link spanning residues 147_272 and DE peptide at residue 272 in the first cluster of ADT2_HUMAN were placed in a separate cluster of ADT2_MOUSE. This demonstrates the ability of clustering to detect similar conformational state(s) of homologues in their respective *H. sapiens* and *M. musculus* compared samples.

Protein 4F2_HUMAN, amino acid transporter heavy chain, has a single intra-protein quantitation cluster with 16 cross-links (including 4 DE peptides) spanning diverse protein lysine residues (Figure 4A). These intra-protein cross-links with similar quantitation among a large variety of sample comparisons most likely have abundance levels that primarily reflect protein levels that originate from a single conformational state. One can assess whether these cross-links are consistent with available PDB structures and AlphaFold models. Interestingly, all cross-links except that spanning residues 245_615 are consistent with most structures. The exceptional cross-link 245_615 has Jwalk SASDs in all structures greater than the maximum for the iqPIR crosslinker, even when allowing for multimeric protein contexts (structure 7DF1 has 4 copies of 4F2_HUMAN). Figure 4B shows the cluster cross-links in the context of structure 2DH2 of the protein. Since the quantitation of this cross-link with an inconsistent SASD of 64 Å is indistinguishable from the other cluster cross-links with consistent SASDs, it is possible that the SASD of this cross-link in the structures does not adequately reflect the protein conformation in samples.

**Figure 4.**
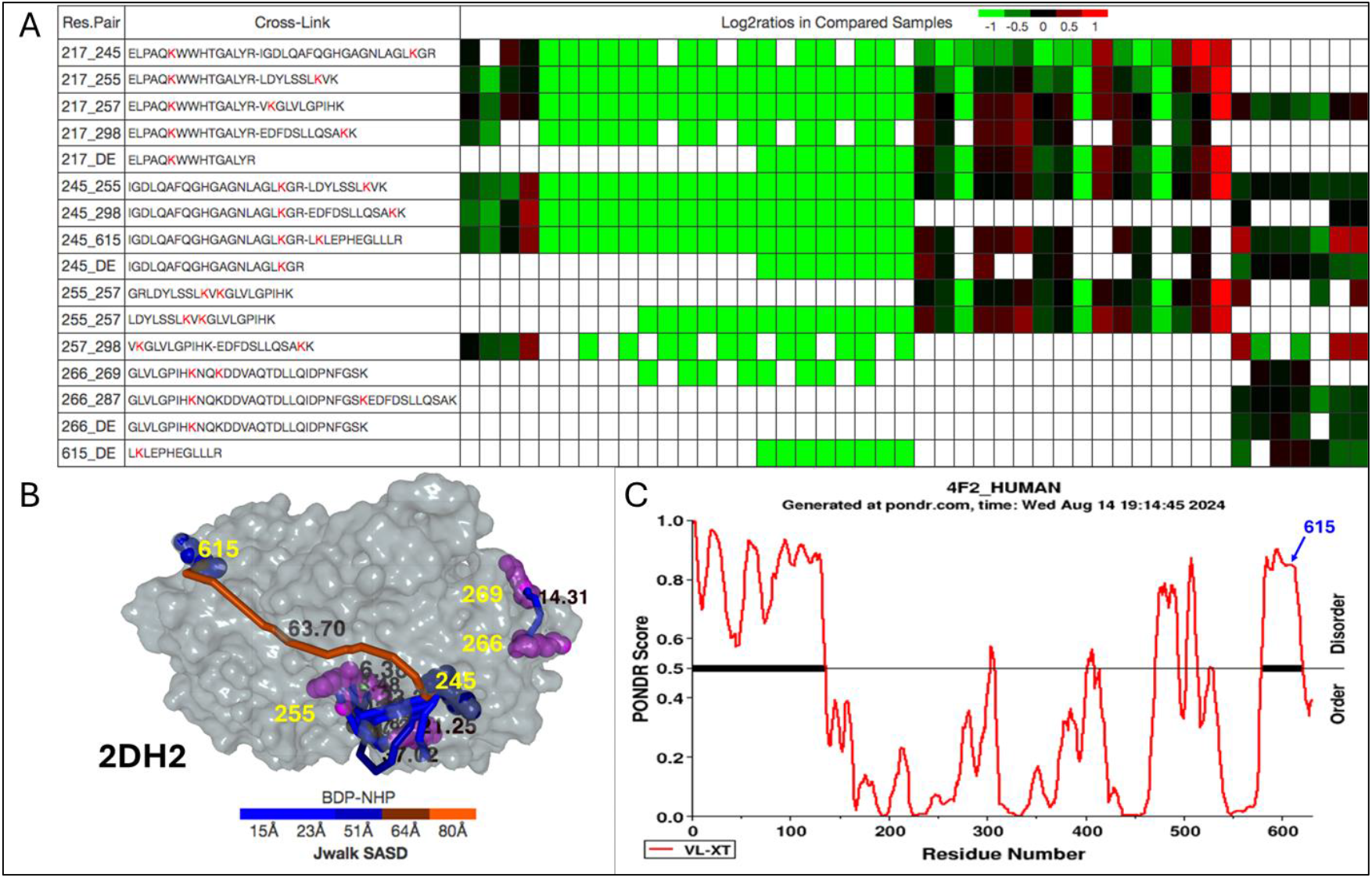
Lone intra-protein cross-link quantitation cluster of 4F2_HUMAN. A. Quantitation of intra-protein cross-links in compared samples. Log2ratios are indicated by color ranging from bright green (−1) to black (0) to bright red (+1) with missing values in white. A residue pair ending in _DE indicates a DE peptide. B. Cross-links in the context of 4F2 structure 2DH2 colored by Jwalk SASD ranging from consistent (blue) to borderline (black) to overlength (orange). DE peptide lysine residues are colored on a similar scale by DSSP accessibility while cross-linked lysine residues are colored purple. The cross-link spanning residues 245_615 has an SASD of 64 Å well in excess of the cross-linker length. C. PONDR analysis of the sequence of 4F2_HUMAN predicting high disorder at residue 615 that may not be well represented in known structures.

We applied PONDR^36^ to predict natural disordered regions of the protein based on sequence and found that residue 615 near the C-terminus is predicted to be highly disordered (Figure 4C). This supports the notion that this cross-link originated from a protein conformation different from known structures, with its C-terminus flexible and within cross-linkable distance of residue 245. Because the quantitation of this and other cross-links are similar, they either originate from a common single conformation with a flexible C-terminus or from two alternative conformations, the known structure and one with a flexible C-terminus, provided they are at fixed relative concentrations in all the compared samples. This would be expected if the C-terminus were within cross-linkable distance of residue 245 a fixed fraction of the time. This highlights that cross-links of a single cluster may originate from multiple conformational states provided those states have fixed relative abundance in all compared samples

The *M. musculus* ACADV_MOUSE protein, mitochondrial very long-chain specific acyl-CoA dehydrogenase, has 2 major quantitation clusters among its intra-protein cross-links (Figure 5). Six of the seven cross-links in the second cluster were reported to have decreased abundance in old versus young mouse muscle^22^. Spanning the binding pocket of fatty acyl-CoA ligand substrate, they were proposed to preferentially originate from protein unbound to substrate and have diminished abundance in old mouse muscle due to increased protein association with fatty acyl-CoA ligand. Since the additional cluster cross-link spanning protein residues 277_483 that wasn’t quantified in that study displayed quantitation similar to the others in all compared samples, it can be inferred that it likely reflects preferential origin from the same conformational state(s) as the other six, *i*.*e*. protein unbound to fatty acyl-CoA ligand, and would be predicted also to have decreased abundance in old versus young mouse muscle. The cross-link spanning similar residues, 279_483 that had diminished abundance in old mouse muscle displayed very similar quantitation to 277_483 in common sample comparisons, with a linear correlation coefficient of 0.84 (Figure S3). This demonstrates how cross-links with similar quantitation that are clustered together comprise an equivalence class that are expected to have similar quantitation in all additional comparable samples.

**Figure 5.**
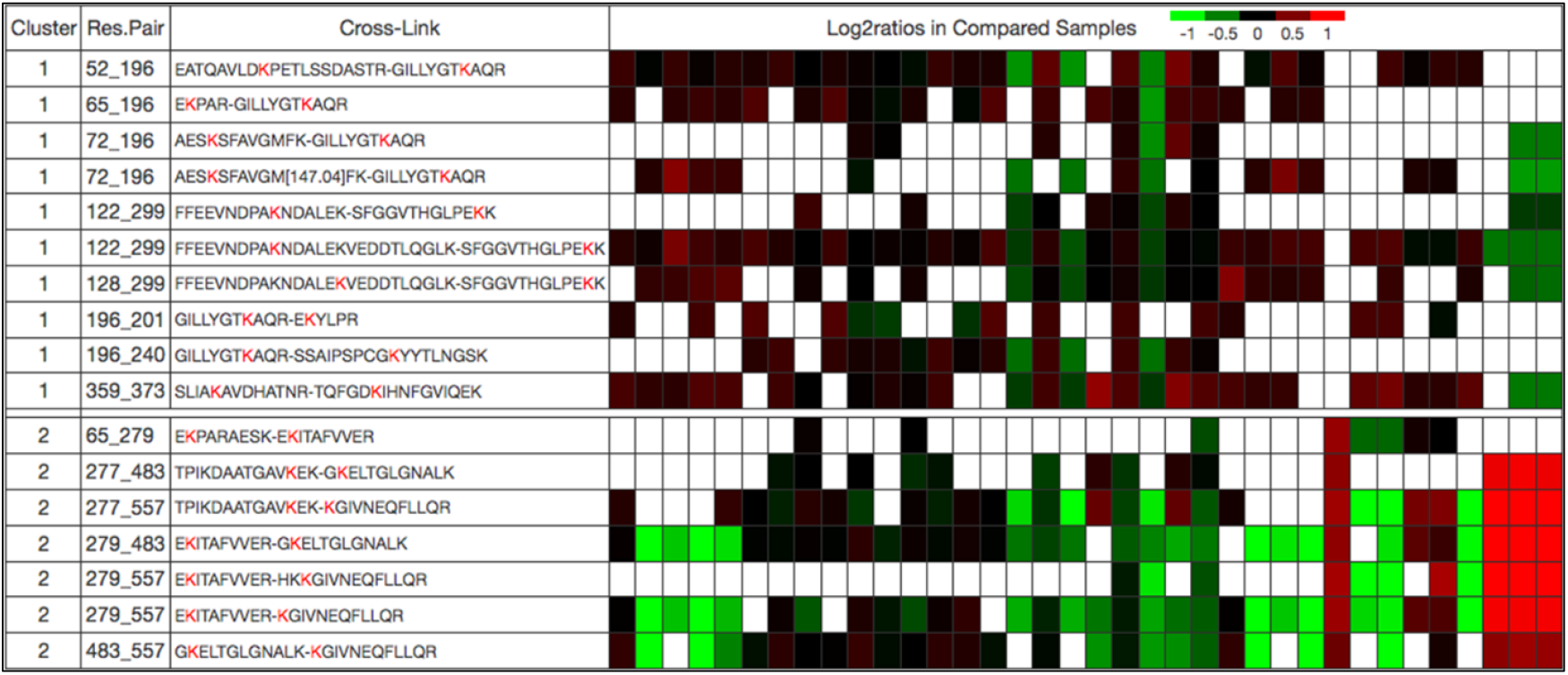
Two intra-protein cross-link quantitation clusters of ACADV_MOUSE. Log2ratios are indicated by color ranging from bright green (−1) to black (0) to bright red (+1) with missing values in white. Cross-links in the second cluster are proposed to originate from a ligand substrate unbound conformational state that is diminished in old versus young mouse muscle and include a cross-link spanning residues 277_483 that was not quantified in that study but presumed based on clustering to have similarly diminished abundance in old versus young mouse muscle.

All nine cross-links in the first cluster were shown to have unchanged abundance in young versus old mouse muscle and were proposed to be insensitive to the binding of substrate ligand fatty acyl-CoA, their quantitation reflecting total protein levels. Thus, the clustering of intra-protein cross-links for this protein across many different compared samples identified two sets of cross-links similar to those described in the qXL-MS study of aged mouse muscle, one with an additional cross-link member. Whereas cross-links in the first cluster are proposed to originate from protein with or without bound fatty acyl-CoA ligand, those in the second cluster predominantly originate from the substrate unbound protein. The conformational influence on these two sets of cross-links applies to a variety of different sample types. Thus, as with ADT, these results based solely on global quantitation were able to identify two sets of cross-links with abundance differentially reflecting multiple protein conformational states.

## Conclusions

Here, we have investigated a new aspect reflecting the value of the growing volume of quantified cross-link data to reveal concerted changes in cross-link levels to better visualize protein plasticity in living systems. The quantitation of cross-links across a wide variety of sample comparisons serves as a signature reflecting encountered protein conformational states. These can include alternative conformations, PTMs, and interacting protein, nucleotide, and ligand partners that affect the likelihood of generating some intra-protein cross-links. Cross-links with similar quantitation can be inferred to have similar relative likelihoods of originating from all encountered protein conformational states with differential relative abundance in compared samples.

The clustering of intra-protein cross-links of a particular protein according to their quantitation in a large number of compared samples enables the grouping of cross-links with respect to the protein conformational state(s) from which they predominantly originate, and thus offers insight into the conformational plasticity of that protein. A single cluster may indicate the presence of only a single protein conformational state, the abundance of which determines the quantitation of all its intra-protein cross-links. However, the existence of multiple clusters indicates intra-protein cross-links with different quantitation that must reflect different relative likelihoods of originating from more than one conformational state present in the compared samples, including alternative conformations, PTMs, and interacting partners. The greater the number of observed clusters, the greater the number of protein conformational states likely present in the samples. We showed that the great majority of proteins analyzed in this study indeed had more than a single cluster indicating protein conformational plasticity. These clustering results are maintained as a publicly available community resource on XLinkDB and increasing numbers of qXL-MS datasets will advance the number of proteins and the ability to resolve clusters of cross-linked peptides.

We demonstrated that clustering by quantitation grouped cross-links of ADT2_HUMAN according to their varying preferences for the m- and c-state protein conformations and grouped cross-links of ACADV_MOUSE according to their proposed sensitivity to protein association with fatty acyl-CoA ligand substrate. We also showed how an intra-protein cross-link of 4F2_HUMAN that has a Jwalk SASD inconsistent with all structures, is nevertheless clustered together with many other intra-protein cross-links that have SASDs that are consistent with those structures. The observation that all cross-links have similar quantitation supports their originating from the same protein conformational state(s). Because the cross-link with an inconsistent SASD is attached to the protein near its C-terminus in a region of predicted high disorder, it is likely that known structures do not adequately represent the protein conformation in the samples or that two conformations coexist in constant relative proportions, one of known structures and one with the C-terminus within cross-linkable distance.

Cross-links clustered together based on having similar quantitation comprise an ‘equivalence class’ with presumed similar relative likelihoods of originating from the encountered protein conformational states that comprise the ensemble in the compared samples. They can thus be considered quantitatively redundant, always expected to have similar quantitation until new samples are analyzed that have altered abundance of new conformational states from which the cross-links have different likelihoods of originating. Interestingly, often multiple cross-links spanning a protein residue pair are identified and quantified. These include cross-linked peptides with oxidized methionine and missed tryptic cleavages. In 80% of such cases the cross-links are clustered together with the majority of other cross-links spanning that residue pair indicating that they often have similar quantitation across many compared samples and provide a testament to the quality of the clustering.

Determining the conformational state(s) from which cross-links of a cluster are derived can be challenging. On XLinkDB, cluster cross-links can be viewed in the context of PDB structures to assess whether they have consistent SASDs and whether they localize to a known binding interface. It may also be beneficial to model alternative protein conformations with software such as AlphaLink^37^ using, in turn, all members of each cluster for guidance. Finally, the clustering of cross-links by quantitation can be extended to inter-protein cross-links spanning a protein pair. Inter-protein cross-link clusters would similarly reflect the levels of the interacting protein conformational states and further advance systems-level quantitative structural biology measurements.

## SUPPORTING INFORMATION

The following supporting information is available free of charge at ACS website

- Figure S1 – Cumulative distribution of DE peptide attached lysine residue DSSP accessibilities in structures.
- Figure S2 – Comparison of intra-protein quantitation clusters of ADT2_HUMAN and those of isoform ADT3_HUMAN and homologue ADT2_MOUSE.
- Figure S3 – Comparison of quantitation log2ratios of ACADV_MOUSE cross-links spanning residues 277_483 and 279_483.
- Video S1 – Training tutorial on how to view intra-protein quantitation clusters for *H. sapiens* and *M. musculus* proteins on XLinkDB.

## Acknowledgements

The authors acknowledge and thank all members of the Bruce lab for helpful comments and suggestions during the course of preparation of this manuscript. This work was supported by the following grants from the National Institutes of Health: R35GM136255, R01HL144778, and R01AG078279.

