## Supplemental Figures for "Large-Scale Quantitative Cross-Linking and Mass Spectrometry Provides New Insight on Protein Conformational Plasticity within Organelles, Cells, and Tissues"

### Supporting Information

**Table of Contents** (list of supplementary components)


- Figure S1 – Cumulative distribution of DE peptide attached lysine residue DSSP accessibilities in structures.
- Figure S2 – Comparison of intra-protein quantitation clusters of ADT2_HUMAN and those of isoform ADT3_HUMAN and homologue ADT2_MOUSE.
- Figure S3 – Comparison of quantitation log2ratios of ACADV_MOUSE cross-links spanning residues 277_483 and 279_483.
- Video S1 – Training tutorial on how to view intra-protein quantitation clusters for *H. sapiens* and *M. musculus* proteins on XLinkDB.

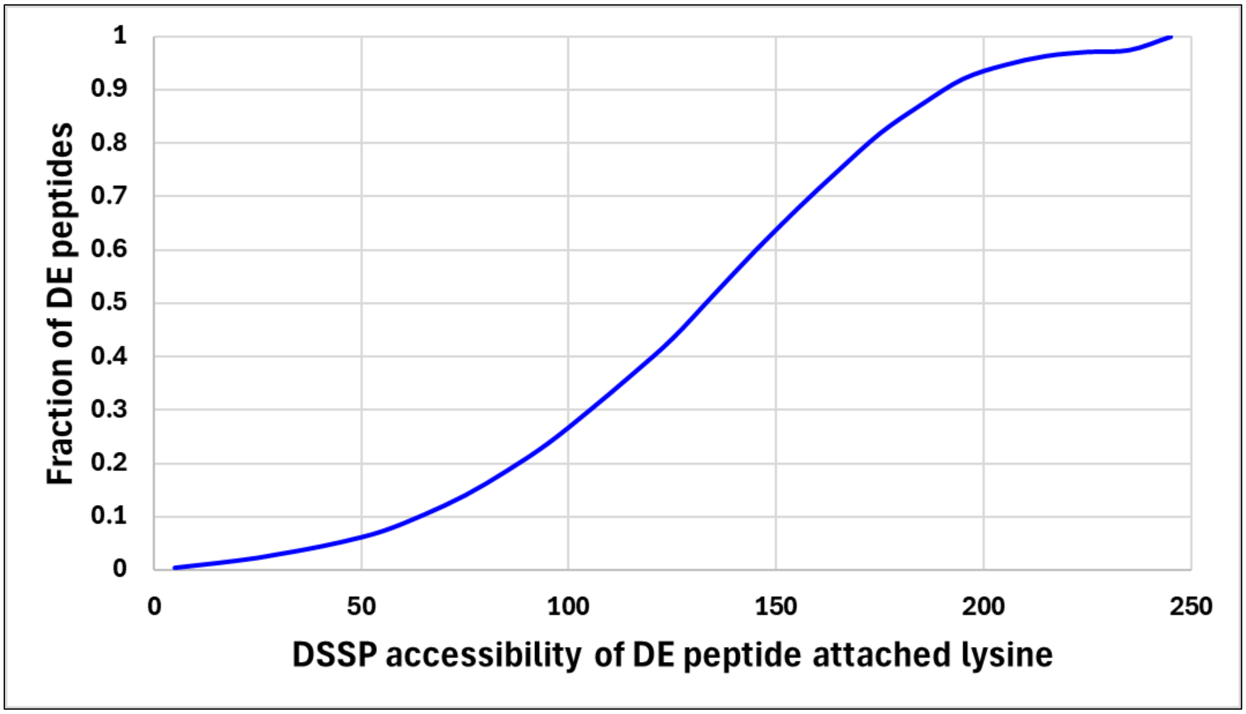


**Supporting Figure S1. Cumulative distribution of DE peptide attached lysine residue DSSP accessibilities in structures.** Shown is the fraction of 9,899 non-redundant DE peptides on XLinkDB with accessibilities in structures less than the indicated value on the x-axis. 95% of DE peptides have an accessibility in a structure of 50 or greater.

| ADT2_HUMAN | ADT3_HUMAN | ADT2_MOUSE |
|---|---|---|
| 23_33 | 23_33 | 23_33 |
| 23_272 | 23_272 | 23_272 |
| 33_147 | 33_147 | 33_147 |
| 33_245 | 33_245 | |
| 33_272 | 33_272 | 33_272 |
| 43_272 | 43_272 | 43_272 |
| 63_163 | 63_163 | |
| 147_272 | 147_272 | 147_272 |
| 163_272 | 163_272 | |
| 272_DE | 272_DE | 272_DE |
| 23_92 | 23_92 | 23_92 |
| 23_94 | 23_94 | 23_94 |
| 23_96 | 23_96 | 23_96 |
| 23_105 | | 23_105 |
| 23_DE | 23_DE | 23_DE |
| 92_DE | 92_DE | 92_DE |
| 92_96 | | 92_96 |
| 92_105 | 92_105 | 92_105 |

**Color Legend**

| |
|---|
| ADT2_HUMAN Cluster 1: m-state |
| ADT2_HUMAN Cluster 2: c-state |
| ADT2_HUMAN Cluster 3: either state |
| ADT3_HUMAN Additional cluster |
| ADT2_MOUSE Additional cluster |

**Supporting Figure S2. Comparison of intra-protein quantitation clusters of ADT2_HUMAN and those of isoform ADT3_HUMAN and homologue ADT2_MOUSE.** Shown are cross-links of ADT2_HUMAN (indicated by residue pair with _DE specifying a DE peptide) that were also assigned to clusters of ADT3_HUMAN and/or ADT2_MOUSE. Assigned clusters for cross-links of the three proteins are indicated by color as equivalent ADT2_HUMAN m-state specific (yellow), c-state specific (green), and state-insensitive (blue), as well as additional clusters of ADT3_HUMAN (pink) and ADT2_MOUSE (red).

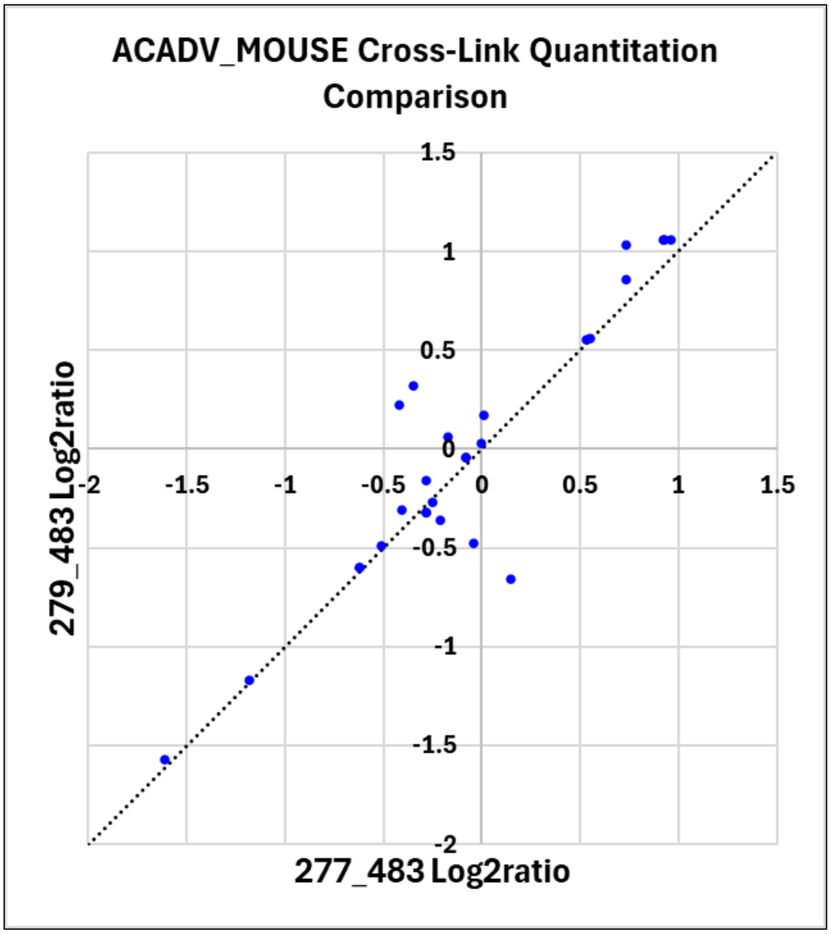


**Supporting Figure S3. Comparison of quantitation log2ratios of ACADV_MOUSE cross-links spanning residues 277_483 and 279_483.** Shown are log2ratios with respect to 27 common compared samples in which they were both confidently quantified. The linear correlation $R^2$ is 0.84. The dashed line indicates equal log2ratios.
